# Benchmarking Nanopore Sequencing for Autosomal and Y-STR Profiling on R10.4.1 Flowcell across Basecalling Models

**DOI:** 10.64898/2026.07.10.737817

**Authors:** Mohammad S. Alsuwaidi, Abdullah Albastaki, Hanan Almulla, Ahmed K. Omar, Mohamed A. Almarri

## Abstract

Short tandem repeat (STR) profiling is the cornerstone of forensic DNA analysis, traditionally performed via capillary electrophoresis. Recently, next generation sequencing has gained prominence due to its increased discriminatory power and enhanced performance with degraded samples. Nanopore sequencing offers a portable and cost-effective alternative, however historically high error rates have precluded its forensic adoption. Here, we evaluate R10.4.1 flow cell chemistry and multiple basecalling tiers (HAC, SUP, HYP) across several iterations (v4.2, v5.0, v5.2, v6.0) to assess their impact on genotyping accuracy. Analyzing 45 STR loci (22 autosomal and 23 Y-STRs) across single-source controls, we introduce a parallelized, user-friendly pipeline designed to transform raw POD5 files into STR profiles. Our results demonstrate a progressive improvement in genotyping accuracy with each basecalling iteration, with the latest models achieving 99.0% autosomal and 100% Y-STR concordance. Furthermore, we find that filtering on raw-read quality scores significantly improves genotyping by reducing background noise and generating cleaner profiles. Notably, the HYPv5.0 Q20 filter drove an average 53.2% reduction in misaligned reads across all loci in comparison to earlier basecalling models. Our study demonstrates that continual bioinformatic improvements in basecalling models, coupled with R10.4.1 chemistry, can provide accurate STR profiles in single-source samples, warranting larger validation studies with more diverse samples to further evaluate performance.

## 1 Introduction

Short tandem repeats (STRs) serve as the foundation of contemporary forensic DNA profiling.[1, 2] Their high polymorphism rate and broad distribution throughout the human genome make them ideal for human identification.[1] Capillary electrophoresis (CE) is the predominant method for DNA profiling, in which loci are PCR-amplified, fluorescently labelled, and subsequently separated by fragment length and dye color.[1, 3, 4]

Although CE is recognized for its reliability and robustness, it faces fundamental technical constraints that impede further progress. The first is limited spectral resolution; as the number of fluorescent dyes increases, the deconvolution of overlapping emission spectra becomes increasingly complex.[1, 5] This has traditionally limited multiplexes to six dyes, though recent advancements have expanded this to eight.[6] A second constraint is the limited spatial resolution, as amplicons labelled with the same dye need to occupy distinct size ranges to avoid overlap. This requires the design of primers which consider the size of amplicons, which can result in larger than ideal fragments, limiting the analysis of degraded samples. Collectively these two constraints limit the number of loci that can be analyzed in a single multiplex reaction.

Next-generation sequencing (NGS) overcomes the limitations of CE-based systems. Rather than resolving alleles based on their fragment length, it determines their nucleotide sequence.[7] Theoretically, tens of thousands of markers, including both STRs and single-nucleotide polymorphisms (SNPs), could be sequenced concurrently, providing discriminatory power several orders of magnitude higher than that of CE systems.[8] Since it does not rely on fragment length separation, smaller amplicons can be designed, improving the analysis of degraded samples. Moreover, NGS can resolve isoalleles, alleles that are identical in fragment length but possess internal sequence variation, which remain indistinguishable using CE methods.[9]

Sequencing platforms such as the Illumina MiSeq FGx and ThermoFisher Ion Torrent have been successfully integrated to forensic workflows, demonstrating that sequence-based STR-profiling is operationally feasible.[10, 11] Although their advantages are very apparent, the high start-up cost and specialized training remain significant barriers for field and forensic laboratories.[7, 8] With the advent of third-generation sequencing, Oxford Nanopore Technologies (ONT) has emerged as potential alternative in the forensic field.[12] Nanopore sequencing determines nucleotide sequences by measuring fluctuations in ionic current as DNA strands translocate through a protein nanopore.[13–15] The MinION platform in particular is compact, portable, and can be deployed in challenging environments.[16, 17] Furthermore, its relatively low acquisition cost makes it accessible to resource-limited laboratories that may otherwise be priced out of NGS technology.[12]

Despite these compelling advantages, nanopore sequencing has historically been hindered by a significant technical limitation: its relatively high raw-read error rate.[18–20] In particular, it is prone to errors in homopolymer regions, stretches of identical nucleotides, which are prevalent in many forensic STR loci.[20] Homopolymers create a constant plateau of electrical signal, making the number of repeated bases ambiguous, resulting in indel errors.[20] While nanopore sequencing is increasingly being established as a mature technology for applications such as metagenomic sequencing and *de novo* assembly, this specific error profile has historically limited its application in forensic testing, where single-base precision is essential for human identification.[21, 22]

Initial investigations into nanopore sequencing for STR and SNP analysis revealed both the promise and the limitations of the approach. These early findings indicated that accurate STR profiling was not yet sufficiently robust for routine forensic applications.[23–26] More recently, the implementation of alignment-based pipelines has enabled the generation of mostly concordant STR profiles from control single-source samples using nanopore data.[27] However, these pioneering studies utilized the R9 flow cell, which has since been discontinued in favour of the R10.4.1 iteration.[27, 28] This newer flow cell introduces several improvements, particularly relevant to forensic STR genotyping. It features a dual-reader pore head that enables the DNA strand to be read twice. This enhances the resolution of homopolymer regions and reduces the indel error rates that hindered previous chemistry iterations.[28, 29] While the newer flow cell improves the physical limitations of its predecessor, the raw current signal generated during translocation must be computationally decoded into a nucleotide sequence, a process known as basecalling. ONT has developed three primary basecalling tiers: FAST, which prioritizes throughput over fidelity; High Accuracy (HAC), which offers a balanced approach; and Super High Accuracy (SUP), which achieves superior accuracy but requires significant computational resources and dedicated Graphics Processing Units (GPUs). These models have undergone several major iterations (most recently versions 4.2, 5.0, 5.2, 6.0), with each successive version demonstrating enhanced accuracy and processing efficiency.

Utilizing the latest R10.4.1 chemistry alongside SUP basecalling, average ONT raw-read accuracy can reach the Q20 (99% accuracy) threshold. Furthermore, ONT recently introduced a fourth tier, the Hyper (HYP) model, currently as a beta release, which provides the highest accuracy to date, albeit with even more demanding hardware requirements. Despite the technical promise of these integrated hardware and software advancements, a comprehensive benchmarking study evaluating their combined performance specifically for forensic STR analysis has yet to be conducted.

In this study, we evaluate the performance of ONT sequencing for profiling a panel of 45 STR loci, consisting of 22 autosomal STRs and 23 Y-STRs. Utilizing single-source control DNA samples sequenced on the R10.4.1 flow cell, we assess genotyping accuracy across HAC, SUP, and HYP basecalling models and their iterations. Complementing this evaluation, we introduce MaSTRspy, a user-friendly pipeline for forensic STR analysis of nanopore data. Inspired by STRspy[27], it adds several features such as a locus-specific genotyping threshold and a graphical user interface (GUI).

## 2 Materials and Methods

### 2.1 DNA Samples and Library Preparation

Three single-source controls (Promega 2800M, Control DNA 007, and NIST C) were amplified using the PowerSeq46GY kit, which targets 45 loci (22 autosomal and 23 Y-STR).[30, 31] Amplification was performed on 1ng of input DNA per sample using 29 PCR cycles, following the manufacturer’s protocol. Sequencing libraries were prepared with the Oxford Nanopore Technologies (ONT) SQK-NBD114.24 ligation sequencing kit, using 200 fmol of amplified product as input. All preparation steps followed ONT protocols without modification. In the first experiment, each control sample was sequenced in triplicate on MinION R10.4.1 flow cells. The second experiment repeated this approach with the same samples using Flongle R10.4.1 flow cells.

### 2.2 Bioinformatics Workflow

Here we introduce our tool “MaSTRspy” available on GitHub (https://github.com/DP-Genome/MaSTRspy) and as a conda package containing all mandatory dependencies. This simplifies installation as it allows users to run the tool without manually editing configuration files. MaSTRspy is a re-written algorithm inspired by STRspy[27], incorporating additional features tailored for forensic STR analysis of nanopore data. The GUI is then run by a simple ‘mastrspy activate’ command.

In this study, raw POD5 files were basecalled and demultiplexed using Dorado v1.0.2. To evaluate the impact of basecalling accuracy on STR genotyping, multiple models were tested, including HAC (versions 4.2, 5.0, 5.2, and 6.0) and SUP (versions 4.2, 5.0, and 5.2) basecalling models. There has not been at the time of this analysis a SUPv6.0 model released by ONT. Furthermore, the POD5 files were also basecalled on the new HYP-5.0 basecalling model. Earlier model versions were also included to quantify improvements in recent ONT basecalling on the same raw POD5 file. We also evaluated filtering reads with a Q-score *>* 20 for the HYP basecalled reads, to evaluate how removing relatively lower quality reads affects mapping and genotyping performance.

We set a minimum read Q-score filter of 10 for the output of each basecalling model and mapped the resulting FASTA to the GRCh38 human reference genome (GCA 000001405.15) using Minimap2 v2.24 [32] using the ONT presets. We note that higher-accuracy models will “rescue” reads that were filtered at Q10 in lower-accuracy models, in particular reads that were near the quality threshold and subsequently reach Q10 with higher models. Mapped reads were then sorted and indexed with SAMtools v1.17 [33] to generate the final BAM files for genotyping against a curated database covering all 45 loci targeted by the PowerSeq 46GY panel (22 autosomal, 23 Y-STR). The database included BED files specifying genomic coordinates on reference genome and FASTA files containing validated allele sequences compiled from both NIST and Hoogenboom datasets to ensure comprehensive variant representation.[9, 34, 35] Following genotyping, MaSTRspy consolidates allele counts into tab-delimited summary files (.tsv) for each sample. STR profiles are visualized using R-based multi-panel plots, where each panel corresponds to a single locus and displays read depth for all mapped alleles.

### 2.3 Analysis Metrics

Within each locus, allele read counts were normalized relative to the most abundant allele. This yields a normalization value of 1.0 for the dominant allele and a value between 0 and 1 for all the remaining alleles.[27] We use different normalization thresholds (0.3, 0.35, 0.4), where a secondary peak is not considered if it lies below this threshold. For Y-STRs we additionally assessed genotype accuracy by choosing the top allele without normalization.

We define “noise” here as reads that map to the incorrect allele, which include stutter positions, for the reference sample divided by the total number of mapped reads at a locus. To assess which STR markers benefit most from better basecalling and quality filtering, we ranked each locus by its noise fraction across all models.

For locus-specific analysis, the noise per sample was calculated, and the average across all samples was determined for each locus. This method ensures equal representation of each sample and prevents any single sample from disproportionately influencing the data. We also calculated stutter percentage at every locus that was either homozygous or had a difference greater than one repeat between the autosomal heterozygous peaks. This allowed us to evaluate the N-1 repeat stutter by dividing its peak height by that of the true allele.

## 3 Results

### 3.1 STR Profile

We first present an example output of our pipeline (Figure 1) which illustrates the allele counts for 45 loci from control DNA 007 basecalled using the SUPv5.2 model. Heterozygous autosomal loci, such as D3S1358 and vWA, exhibited two distinct main allele peaks with similar read depths. In contrast, homozygous autosomal loci such as D13S317 and Y-STR loci, such as DYS19, showed a single dominant-allele peak. We also find reads that map to incorrect reference alleles, though their coverage relative to the main peaks is noticeably lower. For this sample, using the SUPv5.2 basecalling model, all 45 STR loci were recovered and genotyped correctly across all replicates using a normalization threshold of 0.4.

**Fig. 1.**
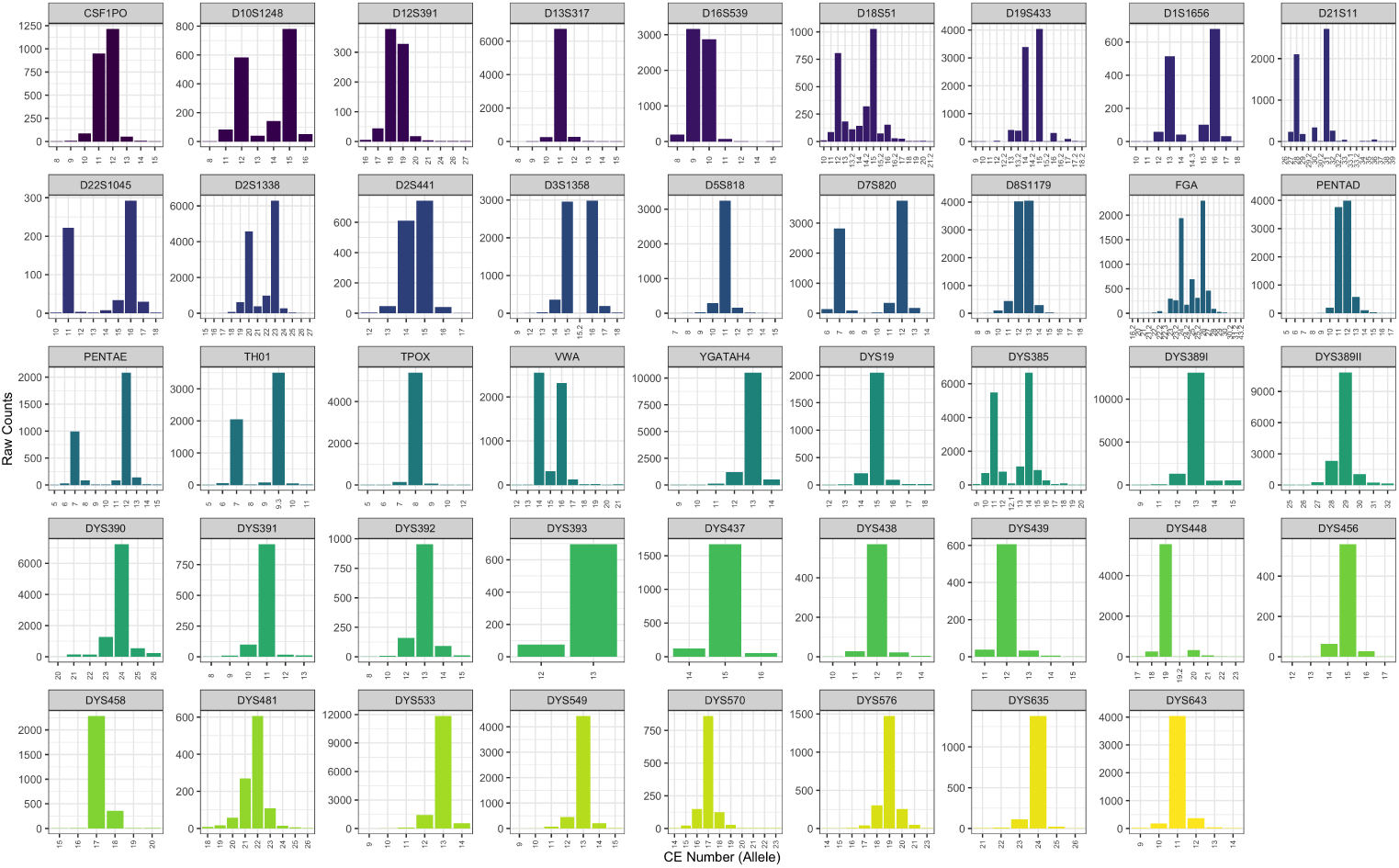
Representative STR profile generated by MaSTRspy for single-source control DNA 007 (barcode 10) basecalled with the SUPv5.2 model. Each panel corresponds to a locus and displays the raw read count (y-axis) for each allele, expressed as the CE allele number (x-axis).

### 3.2 Autosomal Locus Performance

With increased basecalling accuracy from HACv5.2 to SUPv5.2 and HYPv5.0, we generally find that the proportion of reads assigned to the true alleles increased, while noise fractions declined, resulting in a clearer profile at every locus (Figure 2). Figure 3A presents the autosomal STR loci that exhibited the greatest improvement in allele resolution for Control DNA 007 (barcode 10) across the three basecalling models, alongside HYPv5.0 with Q20 filtering.

**Fig. 2.**
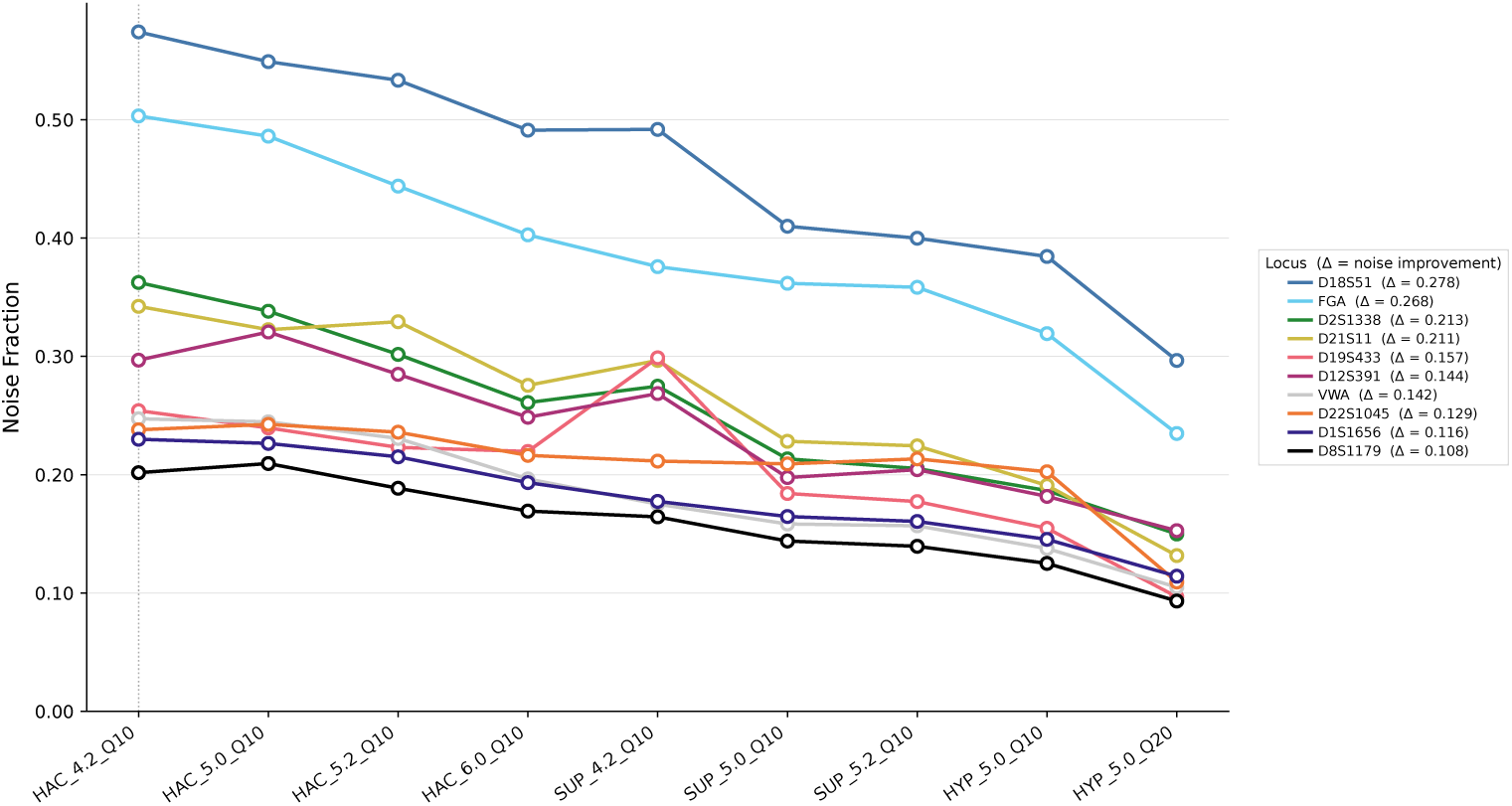
Reduction in autosomal noise fraction with increasing basecalling accuracy. Each line tracks the noise fraction (y-axis) of one autosomal STR locus across successive basecaller models and quality thresholds (x-axis), ordered from lowest to highest accuracy (HACv4.2 Q10 through HYPv5.0 Q20). Noise fraction is the proportion of reads mapping to incorrect alleles. The legend reports each locus’s overall improvement (Δ), defined as its baseline noise fraction (HACv4.2 Q10) minus its lowest value observed across all models.

**Fig. 3.**
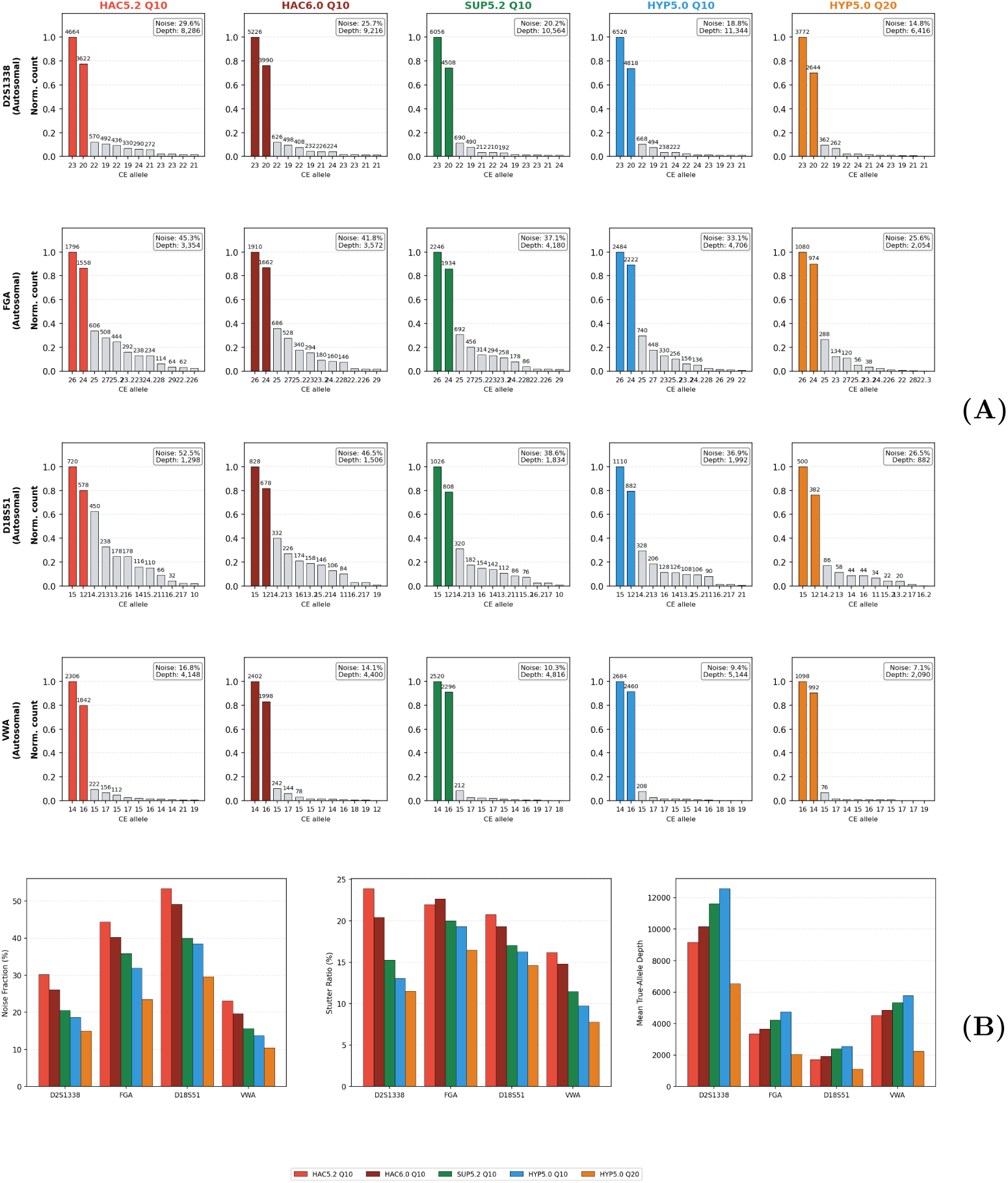
Improvement in autosomal allele resolution with increasing basecalling accuracy. **(A)** Normalized allele profiles for four representative autosomal loci (D2S1338, FGA, D18S51, vWA) in Control DNA 007 (barcode 10). Columns correspond to basecalling conditions of increasing accuracy (HACv5.2 Q10, HACv6.0 Q10, SUPv5.2 Q10, HYPv5.0 Q10, and HYPv5.0 with Q20 filtering); within each panel, bars show the read count for each candidate allele (CE allele number, x-axis) normalized to the most abundant allele (y-axis), with the noise fraction and total read depth annotated. Filled, colored bars represent the true alleles. **(B)** Mean noise fraction, N*−*1 stutter ratio, and mean true-allele read depth for the same loci averaged across all samples, showing the same trend of declining noise and stutter with higher-accuracy models.

Figure 3B shows how each locus performed across all samples. At D18S51, the mean noise fraction dropped from about 53% with HACv5.2 Q10 to 36% with HYPv5.0 Q10, and then to 26.5% with HYPv5.0 Q20. The mean stutter fractions followed a similar pattern, falling from about 21% with HAC v5.2 to around 16%–17% with both SUP v5.2 and HYPv5.0, and down to 15% with HYPv5.0 Q20 filtering. All loci showed similar trends, however some loci showed a more pronounced improvement when filtering for Q20: D18S51, FGA, and D22S1045; which were among the higher-noise loci. Moreover, higher read coverage across all loci was found in basecalling models with higher accuracy as they recovered reads that less accurate models had filtered out at Q10. To help readers explore results for each locus, we have included an interactive HTML viewer that displays these trends across the full loci panel on https://github.com/DP-Genome/MaSTRspy.

Table 1 summarizes concordance between autosomal STR genotypes generated using the basecalling models and the corresponding reference profiles across three normalization thresholds (*>*0.4, *>*0.35, and *>*0.3). The HAC model exhibited systematic errors, such as a false positive at D18S51, that were resolved at the *>*0.4 threshold by the SUP and HYP models. At *>*0.4, SUP v5.0, SUP v5.2, and HYP v5.0 Q10 achieved the highest concordance, while HYP v5.0 Q10 showed a further advantage at lower thresholds, particularly at *>*0.35 and *>*0.3. HYP v5.0 Q20 performed slightly below its Q10 counterpart at *>*0.4 but converged at *>*0.3. At loci with lower coverage, the reduced read depth following Q20 filtering amplifies stochastic allelic imbalance, lowering the normalized height of the true allele below the *>*0.4 threshold and resulting in dropout. Accordingly, the discordances unique to HYP v5.0 Q20 were dropouts at D22S1045 and TH01, rather than drop-ins, indicating that the reduced concordance was driven by diminished coverage rather than increased noise. At the more permissive *>*0.3 threshold, these alleles were recovered, and concordance converged with that of the Q10 filtering. Across all models, the most consistently discordant loci were D10S1248 (dropout), D18S51 (drop-in), and PENTAE (dropout). FGA drop-in errors were observed in HAC and SUP models but were absent in HYP v5.0 Q10. Figure 4 shows the concordance of all samples using latest models at *>*0.4 normalization threshold.

**Fig. 4.**
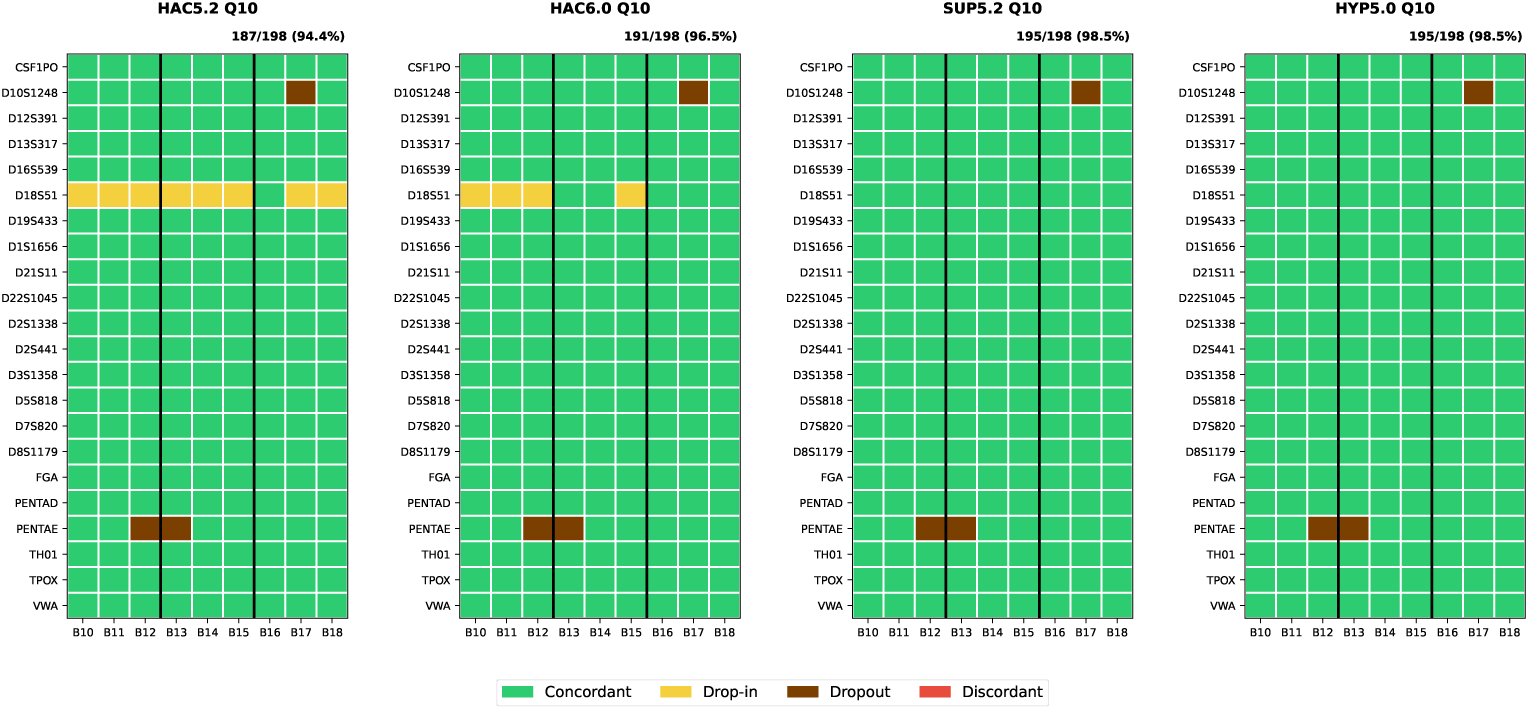
Autosomal STR genotype concordance across all samples at the *>*0.4 normalization threshold. Each panel corresponds to a basecalling model (HACv5.2 Q10, HACv6.0 Q10, SUPv5.2 Q10, and HYPv5.0 Q10); within a panel, each cell shows the genotyping outcome at a locus (y-axis) for a given sample (x-axis), classified as concordant, drop-in, dropout, or discordant, with the overall concordance reported above each panel. The x-axis shows triplicates of the same samples, with B10–B18 indicating barcode 10 to barcode 18.

**Table 1.**
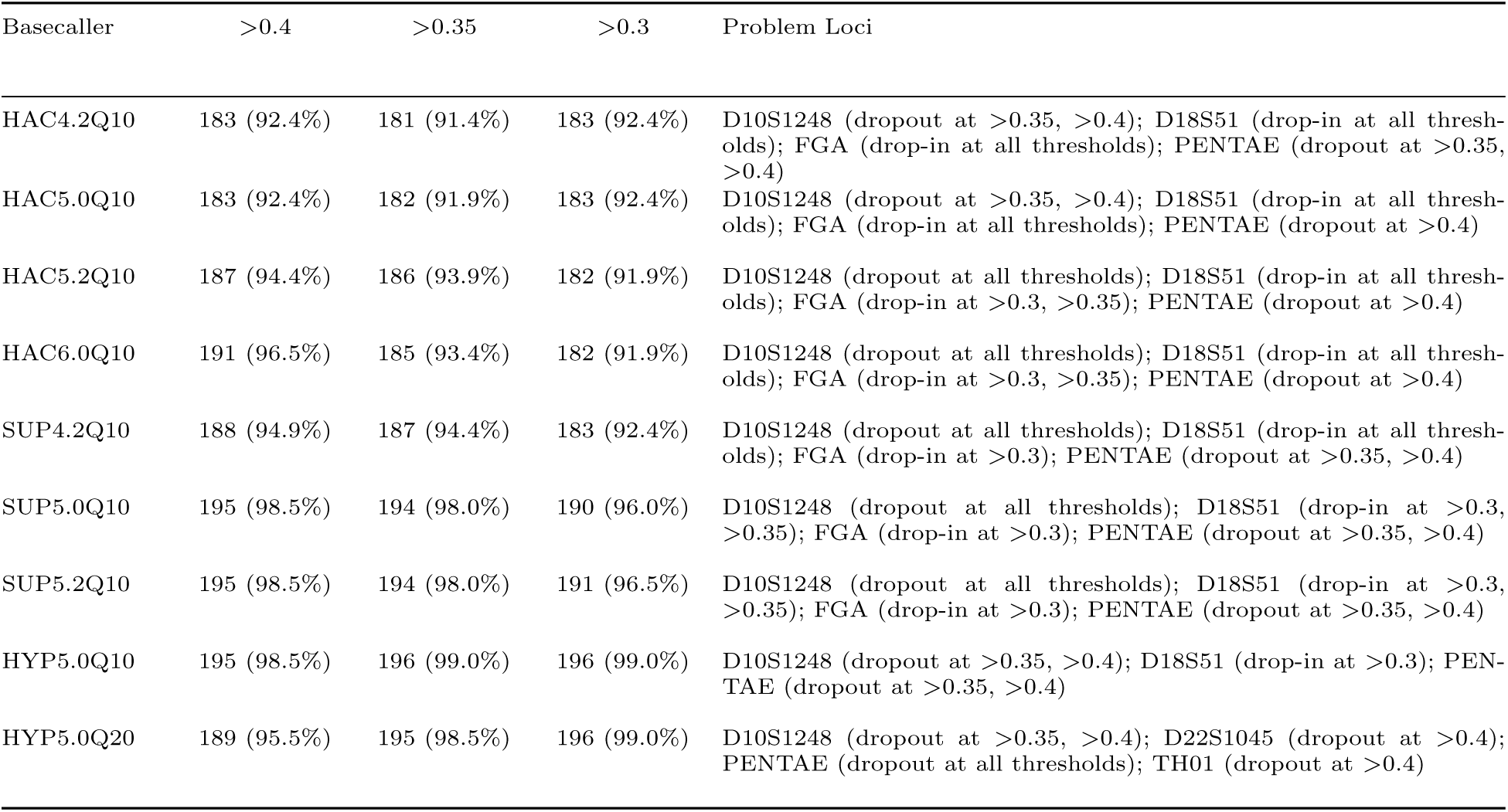
Autosomal loci concordance using various basecallers. Concordance is reported as the number of concordant genotypes out of 198 at each normalization threshold.

### 3.3 Y-STR Per Locus Performance

Similar trends were observed in locus-level improvement in performance across basecalling models for the 23 Y-STR loci (Figure 5). Figure 6A highlights loci that demonstrated the greatest improvement in allele resolution as the accuracy of the basecalling model increased. DYS481, a trinucleotide (CTT) repeat locus, was notable for having substantially higher noise among Y-STRs. At DYS481, the mean noise fraction decreased from 64% with HACv5.2 Q10 to 44% with HYPv5.0 Q10, and further to 39% with HYPv5.0 Q20. Comparable patterns were noted at all other loci including DYS389II, DYS390 and DYS448 where reductions in noise and stutter fractions corresponded with increased reference-allele read depth as model accuracy improved (Figure 6B). We note that some loci show substantially less coverage than others, in particular DYS481.

**Fig. 5.**
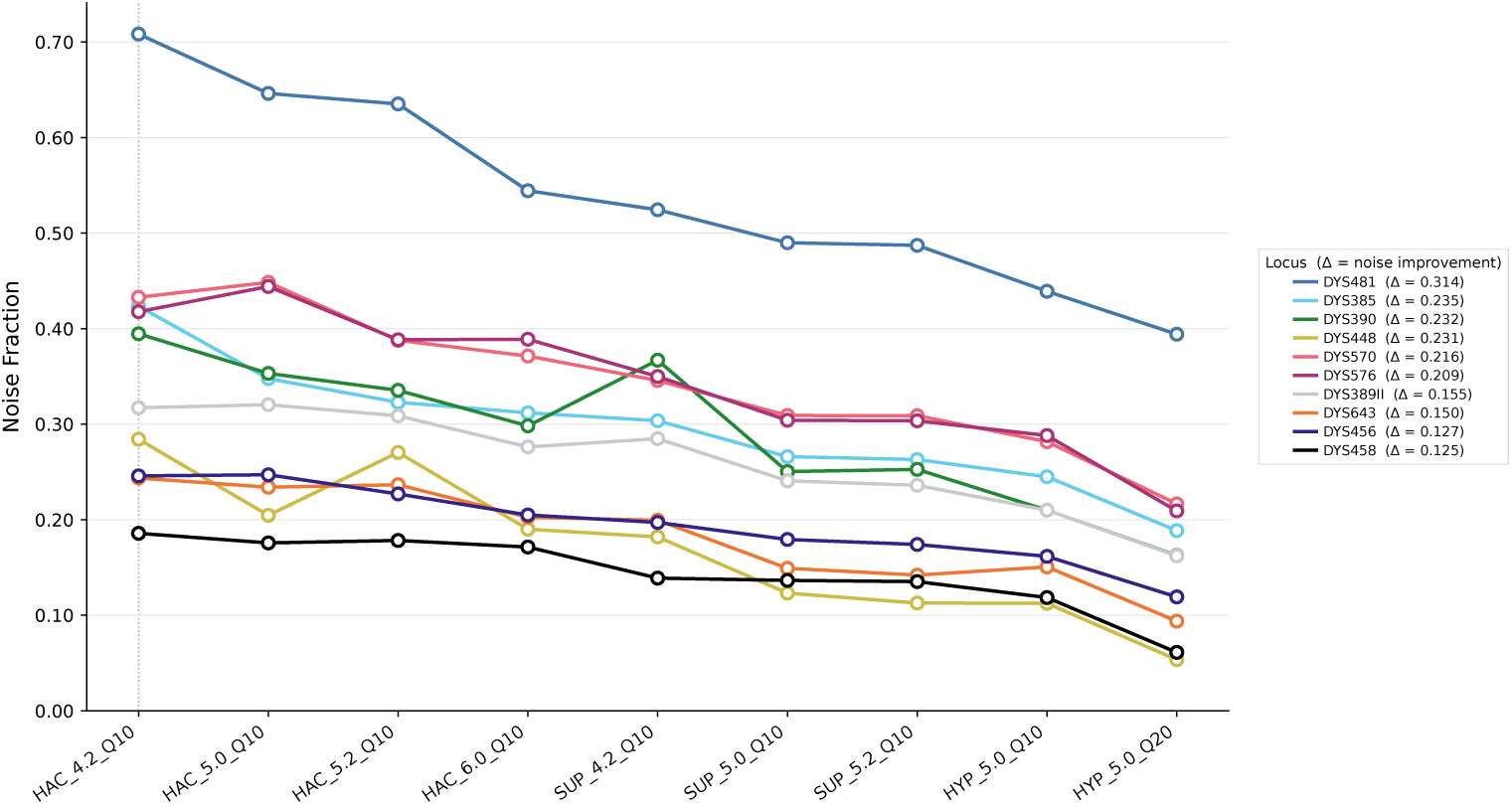
Reduction in Y-STR noise fraction with increasing basecalling accuracy. Each line tracks the noise fraction (y-axis) of one Y-STR locus across successive basecaller models and quality thresholds (x-axis), ordered from lowest to highest accuracy (HACv4.2 Q10 through HYPv5.0 Q20). Noise fraction is the proportion of reads mapping to incorrect alleles. The legend reports each locus’s overall improvement (Δ), defined as its baseline noise fraction (HACv4.2 Q10) minus its lowest value observed across all models.

**Fig. 6.**
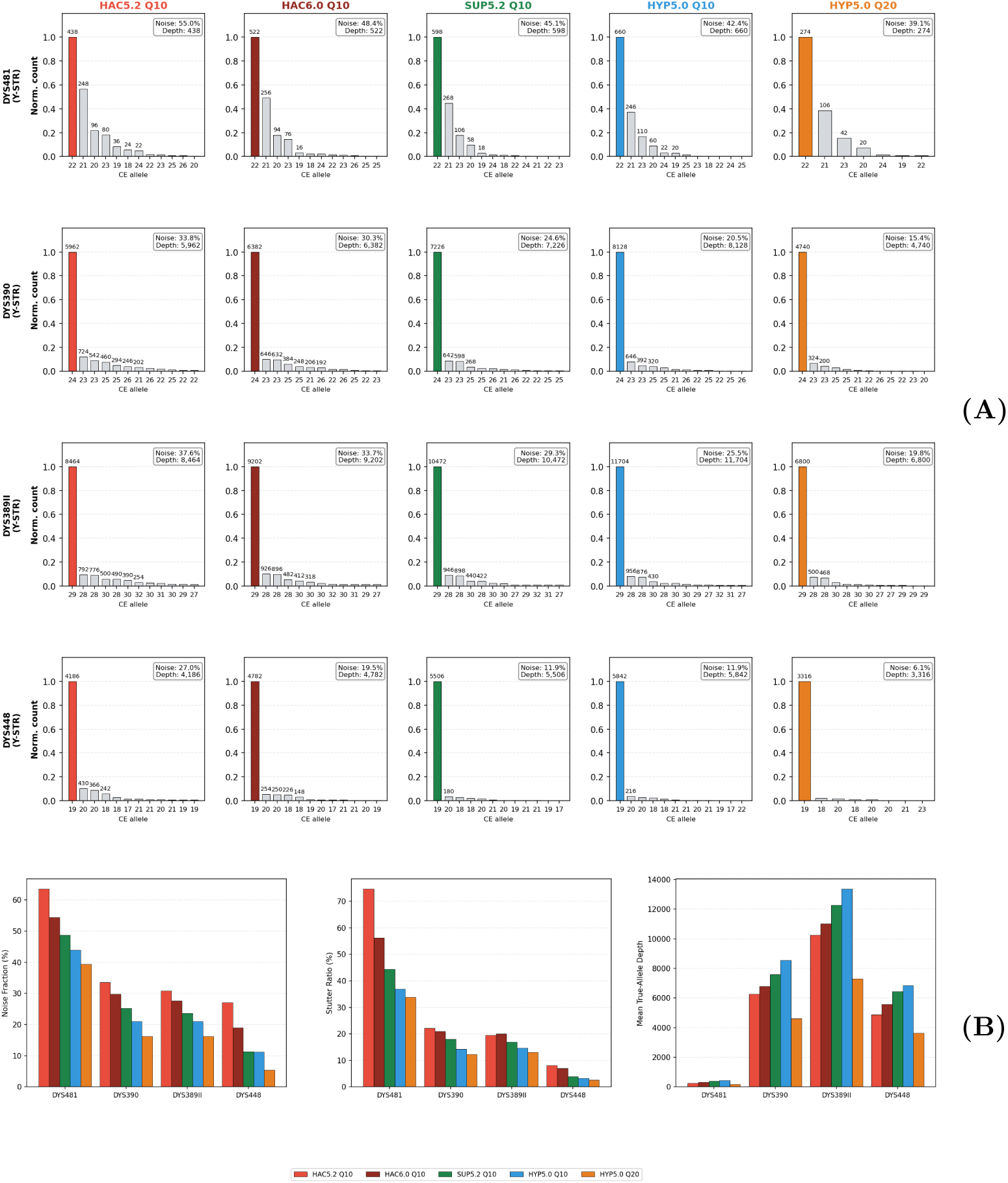
Improvement in Y-STR allele resolution with increasing basecalling accuracy. **(A)** Normalized allele profiles for four representative Y-STR loci (DYS481, DYS390, DYS389II, DYS448) in Control DNA 007 (barcode 10). Columns correspond to basecalling conditions of increasing accuracy (HACv5.2 Q10, HACv6.0 Q10, SUPv5.2 Q10, HYPv5.0 Q10, and HYPv5.0 with Q20 filtering); within each panel, bars show the read count for each candidate allele (CE allele number, x-axis) normalized to the most abundant allele (y-axis), with the noise fraction and total read depth annotated. Filled, colored bars represent the true alleles. **(B)** Mean noise fraction, N*−*1 stutter ratio, and mean true-allele read depth for the same loci averaged across all samples, showing the same trend of declining noise and stutter with higher-accuracy models.

Table 2 summarizes concordance between Y-STR genotypes generated across basecalling models and the corresponding reference profiles at three normalization thresholds (*>*0.4, *>*0.35, and *>*0.3), as well as without normalization by calling the top allele. SUP and HYP models consistently outperformed HAC across all thresholds. At *>*0.4, SUPv5.2 and HYPv5.0 Q10 achieve high concordance, with HYPv5.0 Q20 showing a further improvement at this threshold. Simply choosing the top allele provided 100% concordance at HACv6.0 and all other higher-accuracy models, while older HAC models showed discordances. Figure 7 shows the concordance heatmaps of the top allele for each locus for various basecallers.

**Fig. 7.**
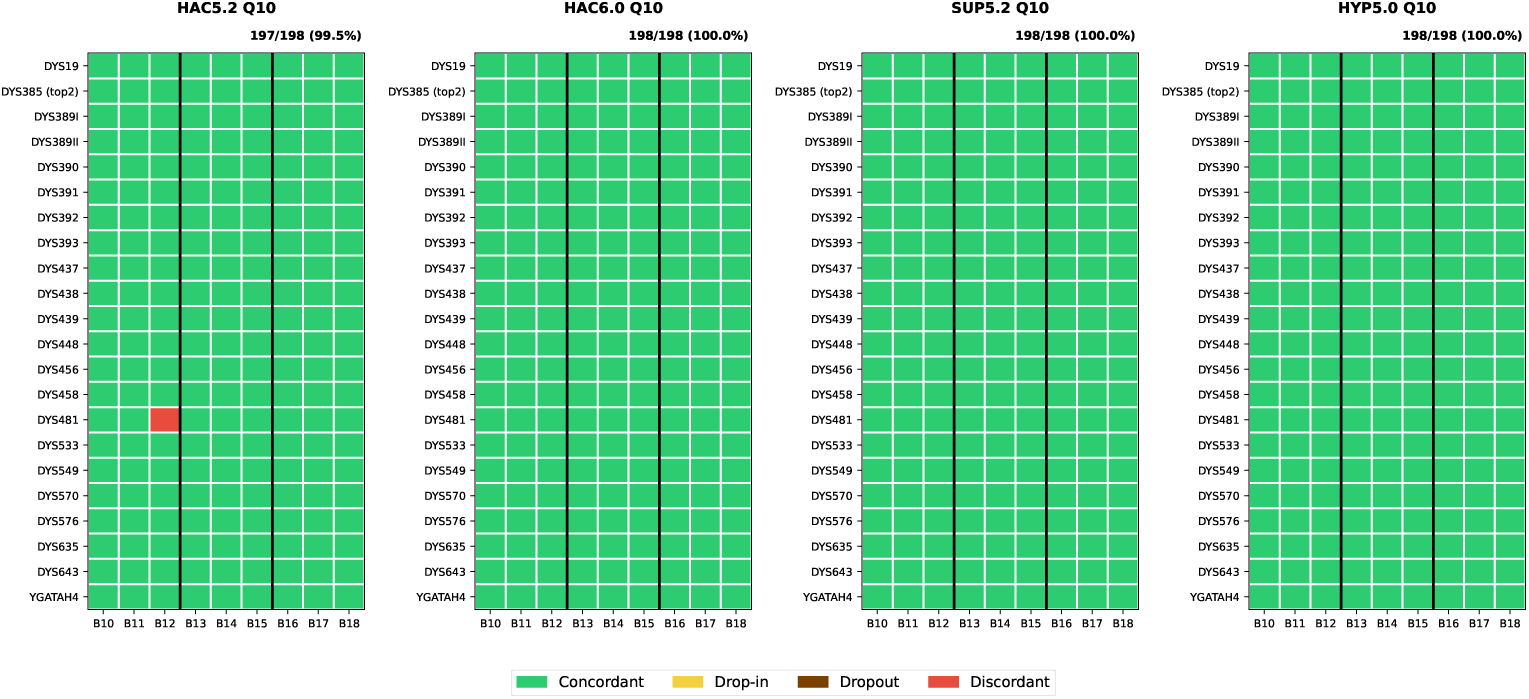
Y-STR genotype concordance across all samples for the top allele called without normalization. Each panel corresponds to a basecalling model (HACv5.2 Q10, HACv6.0 Q10, SUPv5.2 Q10, and HYPv5.0 Q10); within a panel, each cell shows the genotyping outcome at a locus (y-axis) for a given sample (x-axis), classified as concordant, drop-in, dropout, or discordant, with the overall concordance reported above each panel. The x-axis shows triplicates of the same samples, with B10–B18 indicating barcode 10 to barcode 18.

**Table 2.**
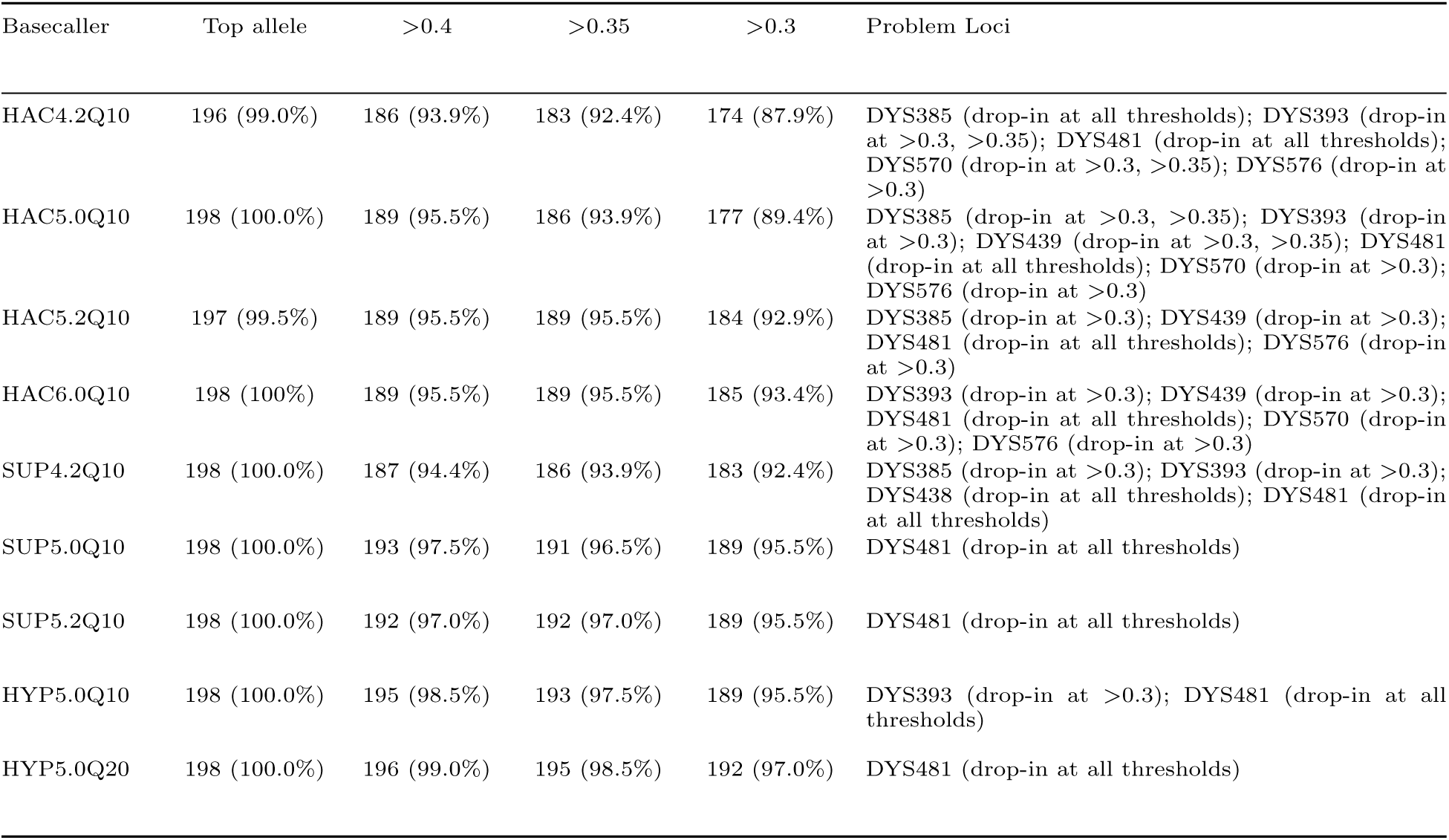
Y-STR loci concordance using various basecallers. Concordance is reported as the number of concordant genotypes out of 198 at each normalization threshold, and for the top allele called without normalization.

| Basecaller | Top allele | >0.4 | >0.35 | >0.3 | Problem Loci |
| --- | --- | --- | --- | --- | --- |
| HAC4.2Q10 | 196 (99.0%) | 186 (93.9%) | 183 (92.4%) | 174 (87.9%) | DYS385 (drop-in at all thresholds); DYS393 (drop-in at >0.3, >0.35); DYS481 (drop-in at all thresholds); DYS570 (drop-in at >0.3, >0.35); DYS576 (drop-in at >0.3) |
| HAC5.0Q10 | 198 (100.0%) | 189 (95.5%) | 186 (93.9%) | 177 (89.4%) | DYS385 (drop-in at >0.3, >0.35); DYS393 (drop-in at >0.3); DYS439 (drop-in at >0.3, >0.35); DYS481 (drop-in at all thresholds); DYS570 (drop-in at >0.3); DYS576 (drop-in at >0.3) |
| HAC5.2Q10 | 197 (99.5%) | 189 (95.5%) | 189 (95.5%) | 184 (92.9%) | DYS385 (drop-in at >0.3); DYS439 (drop-in at >0.3); DYS481 (drop-in at all thresholds); DYS576 (drop-in at >0.3) |
| HAC6.0Q10 | 198 (100%) | 189 (95.5%) | 189 (95.5%) | 185 (93.4%) | DYS393 (drop-in at >0.3); DYS439 (drop-in at >0.3); DYS481 (drop-in at all thresholds); DYS570 (drop-in at >0.3); DYS576 (drop-in at >0.3) |
| SUP4.2Q10 | 198 (100.0%) | 187 (94.4%) | 186 (93.9%) | 183 (92.4%) | DYS385 (drop-in at >0.3); DYS393 (drop-in at >0.3); DYS438 (drop-in at all thresholds); DYS481 (drop-in at all thresholds) |
| SUP5.0Q10 | 198 (100.0%) | 193 (97.5%) | 191 (96.5%) | 189 (95.5%) | DYS481 (drop-in at all thresholds) |
| SUP5.2Q10 | 198 (100.0%) | 192 (97.0%) | 192 (97.0%) | 189 (95.5%) | DYS481 (drop-in at all thresholds) |
| HYP5.0Q10 | 198 (100.0%) | 195 (98.5%) | 193 (97.5%) | 189 (95.5%) | DYS393 (drop-in at >0.3); DYS481 (drop-in at all thresholds) |
| HYP5.0Q20 | 198 (100.0%) | 196 (99.0%) | 195 (98.5%) | 192 (97.0%) | DYS481 (drop-in at all thresholds) |

We further evaluated the effect of Q-score filtering on stutter percentages across Y-STRs, leveraging the hemizygous nature of the Y-chromosomes, using no filtering (Q0), Q10 and Q20. Incremental reduction in mean stutter fractions was observed at all loci as the Q-score filter increased (Figure S1). For instance, mean stutter at DYS576 decreased from 0.179 at Q0 to 0.143 at Q20, and mean stutter at DYS392 declined from 0.160 to 0.136 over the same range. DYS481 exhibited the highest mean stutter among all Y-STR loci, with values of 0.367 at Q0, 0.366 at Q10, and 0.337 at Q20. Q20 filtering was associated with a reduction in mean stutter fraction at this locus; however, stutter values for DYS481 remained higher than those observed for other Y-STR loci. Excluding DYS481, the maximum mean stutter fraction at Q20 was 0.143. Median stutter fraction showed a 20% decrease when Q20 filtering was implemented.

### 3.4 Flongle Performance

The performance of Flongle flow cells was evaluated with SUP v5.2. Figure S2 presents heatmaps of autosomal and Y-STR concordance at the *>*0.4 threshold, alongside Y-STR concordance called by the top allele without normalization. For autosomal STRs, concordance on the Flongle was achieved at 173 out of 198 loci (87.4%), with discrepancies at D10S1248, D12S391, D18S51, D1S1656, D22S1045, FGA, and PENTAE.

For Y-STRs, concordance was observed at 186 out of 198 loci (93.9%) at the *>*0.4 threshold, with discrepancies at DYS393, DYS456, DYS458, DYS481, DYS570, and DYS576, and rose to 197 out of 198 loci (99.5%) when calling the top allele, with the single discrepancy at DYS481.

## 4 Discussion

Our analysis demonstrates that with the current MinION 10.4.1 flowcells and latest basecalling models, nanopore sequencing can generate relatively accurate STR profiles for single source samples. We find that the choice of basecalling model is an important factor influencing genotyping success. Notably, since we used a read alignment-based method to call alleles, improved accuracy by basecalling models of the same raw reads appears to provide better alignment to reference alleles which improved genotyping. However we find that this increase in accuracy is variable across loci, with some loci showing a more pronounced improvement in alignment relative to others.

Despite advancements in basecalling, certain loci show errors due to inherent biological or amplification factors. In particular, D18S51, FGA and DYS481 show consistently elevated noise across all basecallers. The first two are composed mostly of AGAA repeats, which can potentially explain its struggles in ONT sequencing. While the latter is a CTT trinucleotide repeat known to be a high stutter locus.[36] We also find dropouts in PENTAE, however this locus appears to have a low error rate in our analysis, suggesting high allelic-imbalance, rather an intrinsic issue with nanopore sequencing. These dropouts align with a previous study which identified an allele imbalance at this locus when using the PowerSeq 46GY kit.[31] We also show that small differences in normalization thresholds (0.3, 0.35, 0.40) provide different genotyping results. Consequently, we recommend that laboratories adopt locus-specific stutter or normalization thresholds rather than universal ones we apply here.

We evaluated setting a high Q-score (Q20) threshold on genotyping accuracy performance. When used with the HYP5.0 model, Q20 filtering produced a substantially cleaner signal with a marked reduction in noise, enabling superior allele resolution. However, this improvement comes with a significant trade-off: with the HYP5.0 model, often more than half of the total reads were filtered at this threshold, illustrating the current quality-score limitations of nanopore sequencing. This necessitates an internal evaluation balancing Q-score filters with resulting read depth to ensure allele assignments estimates remain reliable. We also assessed the performance on the Flongle flowcell, an inexpensive alternative to the MinION, which showed substantially higher errors. We note that the Flongle has been recently discontinued by ONT.

Adopting the ONT platform in a routine forensic setting introduces unique challenges. The rapid release cycle of ONT basecalling models necessitates frequent internal validations. To mitigate this burden, we propose a “re-analysis” workflow: since only the raw signal POD5 files need to be re-processed through newer basecalling models, laboratories can potentially avoid the sample preparation and sequencing phases of validation for new software releases, provided the raw signal data is stored long-term. However, it is important to note that the high-performing SUP and HYP models require investment in GPU-based computing units. Standard CPUs struggle to process these models within realistic forensic timeframes, representing an additional infrastructure cost that laboratories must factor into their adoption strategy. We also note that the production-ready basecaller version that is installed on the ONT sequencer is always older than the latest version available on GitHub, which requires rebasecalling the raw POD5 using the latest models on a separate workstation.

While our results are promising for single-source samples at optimal concentrations, several hurdles remain for operational casework. Forensic samples are frequently degraded, low-template, or complex mixtures. As noted in a recent ONT-based single-source sequencing study,[37] unexplained genotyping errors at STRs can still occur in forensic samples, although it used the now discontinued 9.4.1 flow cell technology. Future research must involve comprehensive mixture studies varying contributor ratios and DNA quantities. Furthermore, moving away from fixed universal thresholds toward adaptive or probabilistic frameworks, which could be integrated into future iterations of our pipeline, will be essential for handling the complexities of real-world forensic evidence.

## Data Availability

The MaSTRspy pipeline is available on GitHub (https://github.com/DP-Genome/MaSTRspy). The POD5 files used in this study are available on Zenodo (10.5281/zenodo.19954156).

## Acknowledgements

We thank Yasmeen Qutub (Oxford Nanopore Technologies) for technical assistance. The authors of this paper are supported by the General Department of Forensic Science and Criminology, Dubai Police HQ.

## Author Contributions

M.S.A. performed the data analysis, developed the MaS-TRspy pipeline, and co-wrote the manuscript. A.B. and H.A. designed the laboratory experiments and carried out the sample preparation and sequencing. A.K. set up the sequencing and computing infrastructure. M.A.A. designed and supervised the project and co-wrote the manuscript.

## Supplementary Information

**Fig. S1.**
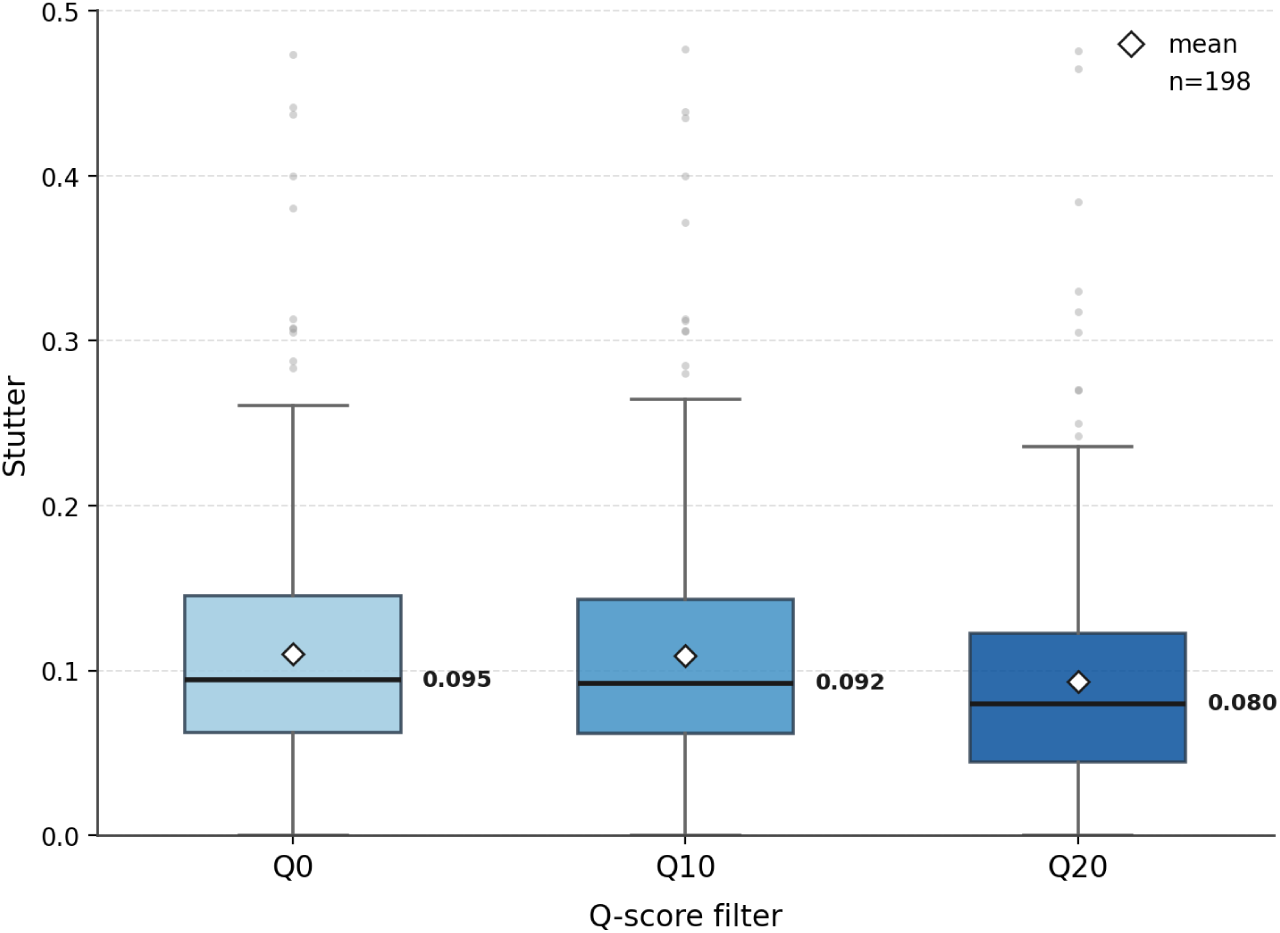
Effect of read Q-score filtering on Y-STR stutter fractions. Box plots show the distribution of stutter fractions across all 23 Y-STR loci (*n* = 198 genotypes) for the HYPv5.0 basecalled reads under three quality-score filtering conditions: no filtering (Q0), Q10, and Q20. In each box, the horizontal line marks the median. The median stutter fraction decreased from 0.095 at Q0 to 0.092 at Q10 and 0.080 at Q20.

**Fig. S2.**
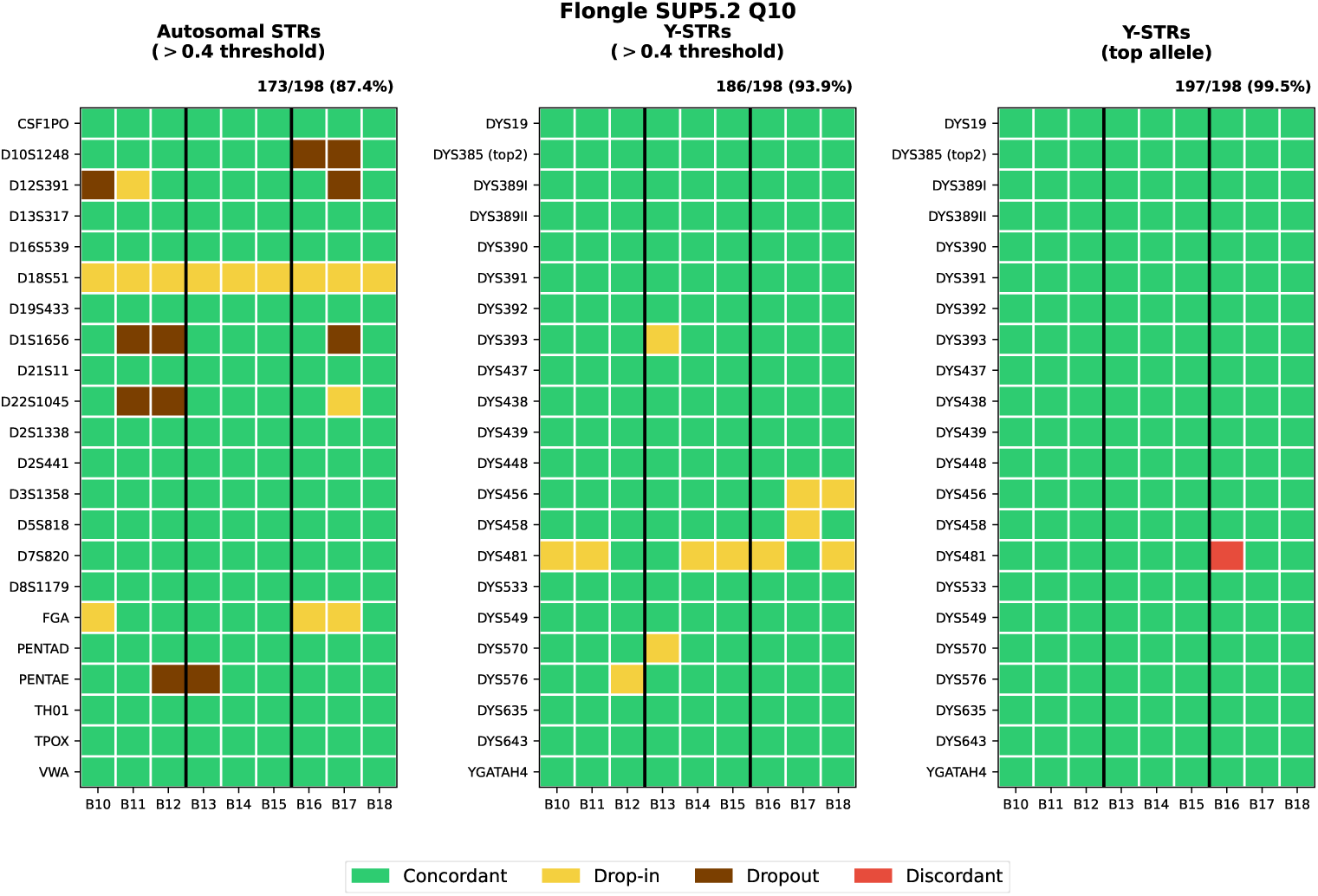
Genotype concordance on the Flongle flow cell. Concordance for the Flongle SUPv5.2 Q10 run shown for autosomal STRs at the *>*0.4 normalization threshold (left), Y-STRs at the *>*0.4 normalization threshold (middle), and Y-STRs called by the top allele without normalization (right). Within each panel, each cell shows the genotyping outcome at a locus (y-axis) for a given sample (x-axis), classified as concordant, drop-in, dropout, or discordant; the x-axis shows triplicates of the same samples, with B10–B18 indicating barcode 10 to barcode 18. Overall concordance is reported above each panel (173/198, 87.4% for autosomal STRs; 186/198, 93.9% for Y-STRs at *>*0.4; 197/198, 99.5% for Y-STRs by top allele).

